# Cerebrospinal fluids from healthy pregnant women does not harbor a detectable microbial community

**DOI:** 10.1101/2020.09.16.299065

**Authors:** Yongyong Kang, Xinchao Ji, Li Guo, Han Xia, Xiaofei Yang, Zhen Xie, Xiaodan Shi, Rui Wu, Dongyun Feng, Chen Wang, Min Chen, Wenliang Zhang, Hong Wei, Yuanlin Guan, Kai Ye, Gang Zhao

## Abstract

Cerebrospinal fluids circulating human central nervous system have long been considered aseptic in healthy individuals, because normally the blood-brain barrier protects against microbial invasions. However, this dogma has been questioned by several reports that microbes were identified in human brains, raising the question whether a microbial community is present in cerebrospinal fluids of healthy individuals without neurological diseases. Here, we collected and analyzed metagenomic and metatranscriptomic sequencing data of cerebrospinal fluid specimens from a cohort of 23 pregnant women aged between 23 and 40 and one-to-one matched contamination controls. From data analysis of 116 specimens of eight different types, we detected 619 nonredundant microbial taxa which were dominated by bacteria (75%) and viruses (24%). In cerebrospinal fluids metagenomic samples, a total of 76 redundant species were detected including four (one nonredundant) eukaryota taxa, eleven (four nonredundant) bacteria, and 61 (21 nonredundant) viruses that were mostly bacteriophages. Metagenomic data analysis found no significant difference between cerebrospinal fluid specimens and negative controls in terms of microbial species diversity. In addition, no active or viable microbiome were present in the cerebrospinal fluid samples after subtracting microbes detected in contamination controls. In conclusion, we found no strong evidence that colonized microbial community exist in the cerebrospinal fluids of healthy individuals.

**IMPORTANCE:** Microbiome are prevalent throughout human bodies with profound health implications. However, it remains unclear whether a microbiome is present and active in human cerebrospinal fluids that are long considered aseptic given the blood-brain barrier. Here, we applied unbiased metagenomic and metatranscriptomic sequencing to detect microbiome in cerebrospinal fluids collected from a cohort of 23 pregnant women with matched controls. By analyzing 116 specimens of eight types, no strong evidence was found to support a presence of colonized microbiome in the cerebrospinal fluids. Our findings have profound implications to human immunity against neurological infections and disorders, providing a guide for disease diagnostics, prevention and therapeutics in clinical settings.

## INTRODUCTION

First defined by Joshua Lederberg in 2001(1), human microbiome has since been discovered at almost every part of human bodies such as gut, oral, skin, bladder, vagina, lungs(2–8). It has profound impact on human health, being associated with a broad range of human diseases including cancers, diabetes, schizophrenia and autoimmune diseases etc.(9–12). However, due to difficulties in the identification and traceability of contaminations, it remains controversial whether there are colonized microbial community at some sites such as placenta, blood and amniotic fluid (13–17), although recently both experimental and analytical methods have improved in sensitivity and accuracy regarding microbiome discovery.

Cerebrospinal fluid (CSF) circulating the human central nervous system (CNS) has long been considered sterile given that the blood-brain barrier is thought to effectively protect against microbial invasions. However, this traditional knowledge of microbe-free CSF has been challenged in recent years with several reports of microbes detected in human brains and CSF. For example, a bacterial pathogen *Porphyromonas gingivalis* was identified in brain regions including cerebral cortex and hippocampus in patients with Alzheimer’s disease(18). In addition, a number of DNA viruses in CSF were identified from a subjects of virome(19). It remains elusive whether these reports are evidence of a common microbiome in human CSF and CNS, or simply sporadic and accidental events.

Given the debate over the existence of any microbial community in CSF and the importance of understanding microbial infection in human central nervous systems, we performed microbiome analysis to characterize bacteria, archaea, eukaryota and viruses of CSF from a cohort of 23 donors without neurological disorders, as well as one-to-one matched positive controls (oral and skin) and negative controls (normal saline). DNA/RNA extraction buffers and sterile swab were also collected as controls. In total, 116 specimens of eight types were used in this study. Considering the limitations of 16s ribosomal RNA based approach to achieve a consistent result in species or strain level(20–23), we choose unbiased metagenomic next-generation sequencing (mNGS) for rapidly detecting all genetic materials of microbiome at species resolution and use metatranscriptomic sequencing to assess the physiological states of microbial communities in CSF(24, 25). mNGS as a promising approach, its clinical diagnostic performance in infectious diseases has been widely adopted in the medical community by multi-center studies(26–28).

In our data analysis, we found no significant difference between CSF specimens and negative controls in microbial species diversity. In all CSF samples, no active or viable microbiome was present after subtracting microbial taxa detected in CSF by those detected in contamination controls. Taken together, no strong evidence was found in our study supporting that colonized microbiome exists in the cerebrospinal fluids.

## RESULTS

### Metagenomic sequencing of cerebrospinal fluids in healthy individuals

To investigate whether there is microbiome in CSF, we collected and analyzed microbiome of CSF samples from 23 pregnant women aged 23–40 years who underwent intraspinal anesthesia before the caesarean section via lumbar puncture, coupled with normal saline collected with syringe as negative controls. For each subject, oral and skin microbiomes were also collected and analyzed as positive controls (Figure 1a). All samples were then subjected to DNA extraction and metagenomic shotgun sequencing and analysis. Finally, to validate whether the microbiome, if any detected in CSF, is physiologically active, metatranscriptome profilings for 12 of the pregnant women CSF samples were performed. After quality control (QC) with KneadData(29) (v0.7.2) for metagenomic and metatranscriptomic sequencing data, MetaPhlAn(30) (latest version 3), a state-of-the-art taxonomic classification tool based on unique clade-specific marker genes, was used to detect microbes in each sample.

**Figure 1:**
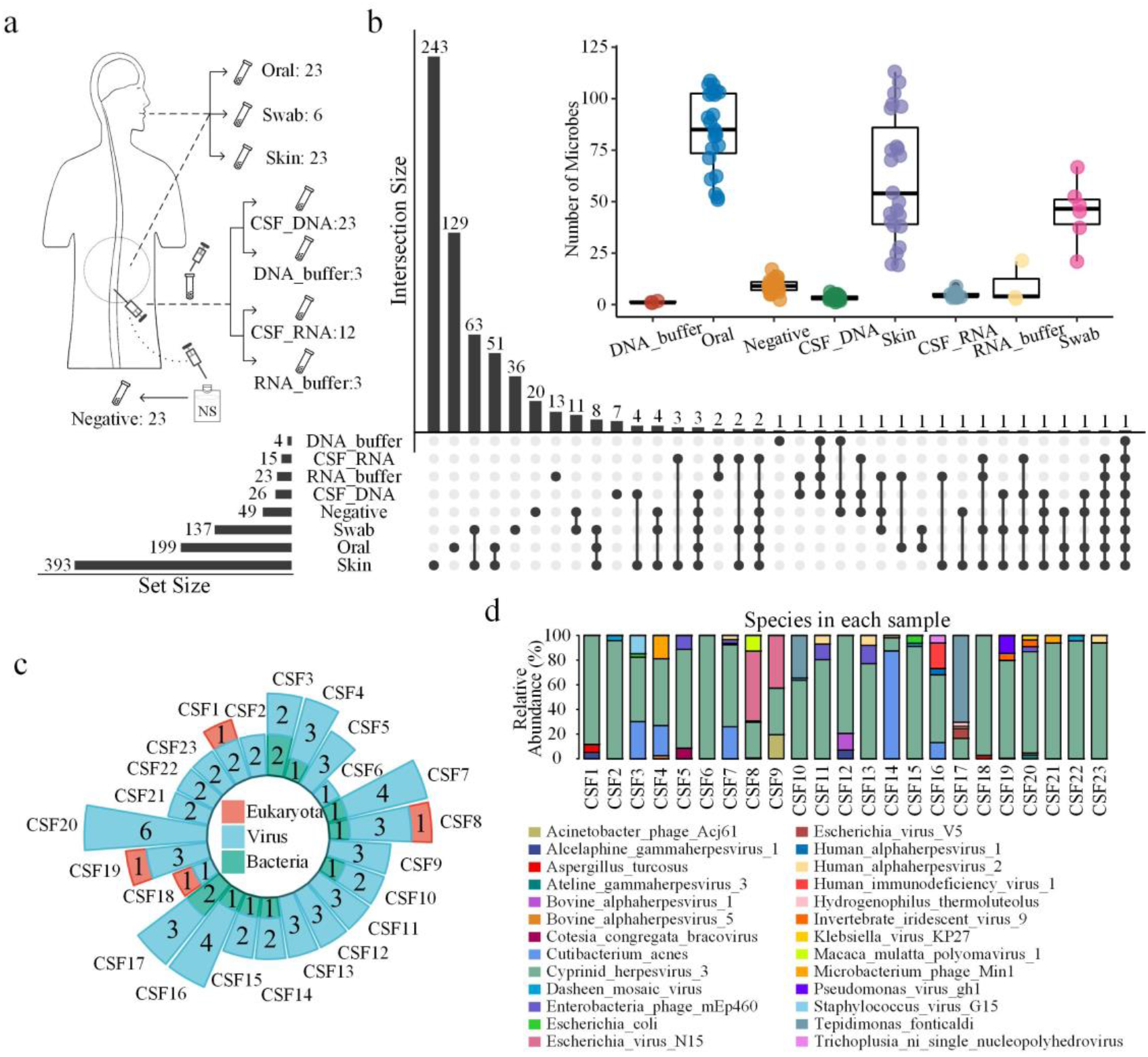
Microbial community structure in human CSF of 23 healthy individuals. (a) Metagenomic experimental design in this study: CSF and matched control samples (positive controls: oral and skin; negative controls: saline solution) collected from 23 pregnant women along with DNA and RNA extraction buffers (number indicates replicates) and were sequenced for metagenomic and metatranscriptomic analysis (see Methods). (b) An overview of microbes detected in each sample type. The number of microbes detected in each sample, and shared species between different samples were shown in the upset plot, with the dots representing intersections among sample types, and the bars representing the number of microbes for each sample type (horizontal bars) and ones shared for each intersection type (vertical bars). The inlet shows a box plot summarizing distributions of the number of species detected for different sample types. (c) Circle barplot summarizing the the number of microbial species in each CSF_DNA sample, categorized into three major types: eukaryota, virus and bacteria. (d) Microbial community structures of 23 CSF_DNA samples shown in a stacked barplot that summarizes the relative abundance of different species of microbes detected for each CSF_DNA sample.

In total, we detected 619 nonredundant microbial taxa in 116 specimens using metagenomic and metatranscriptomic sequencing and analysis (Supplementary Table S1). These microbes detected in all samples were dominated by bacteria (75%) and viruses (24%). Overall, skin, oral and swab samples had the most abundant microbiome of all samples with 393, 199 and 137 nonredundant microbial taxa, respectively. This came as no surprise because skins and orals are well known to harbor a plethora of microbes. By contrast, the number of microbial taxa detected in CSF_DNA (26), negative controls (49) and extraction buffers (27) were relatively fewer (Figure 1b). We then compared the taxa detected in different specimen types, finding that there was little overlap among all samples. Skin, oral and sterile swab had a large amount of unique microbial taxa among all sample types, with 243, 129 and 36 taxa found only in these samples, respectively (Figure 1b). Although a large variation in the number of microbes detected was observed for skins, orals and swabs, a much smaller variation was found for CSF_DNA, CSF metatranscriptomic (CSF_RNA), negative controls and extraction buffers (DNA/RNA buffer) (Figure 1b). The oral samples were rich in *Streptococcus*, *Veillonella*, *Neisseria, Rothia* and *Prevotella,* while the skin samples were rich in C*utibacterium*, *Staphylococcus*, *Micrococcus*, *Malassezia,* consistent with many previous studies(4, 6, 31) (Supplementary Figure 1). Our successful detection of known microbiome for orals and skins provided a proof-of-concept of NGS-based metagenomic sequencing method, laying a solid foundation for our exploration of CSF microbiome using such a method.

We next focused on examining the microbial species detected for each CSF_DNA specimen. In CSF samples, a total of 76 redundant species including 11 (4 nonredundant) bacteria, 61 (21nonredundant) viruses and four (one nonredundant) eukaryota taxa were detected (Figure 1c). Most of these viruses were bacteriophages. The relative abundance of microbes suggested the species “*Cyprinid_herpesvirus_3*” are the predominant species in 19 of 23 CSF_DNA samples (Figure 1d). *Cutibacterium_acnes* in species level appeared in 5 specimens. Additionaly, 100%, 26%, 22%, and 22% of all CSF_DNA samples contain *Cyprinid herpesvirus 3*, *Human alphaherpesvirus 2*, *Enterobacteria phage mEp460* and *Dasheen mosaic virus*, respectively. However, “*Cyprinid herpesvirus 3*” detected in all CSF_DNA samples were also found in all negative and skin samples, suggesting a likely external source of these microbes during the CSF sampling procedure.

### The microbiome signature of cerebrospinal fluids and negative controls is similar

Since microbial species were identified in both CSF_DNA and negative controls, it is likely that microbial cells and/or DNA present in negative controls may have been introduced into CSF during the sampling process. Similarly, the possibility of skin microbiome being introduced into CSF during lumbar puncture could not be ruled out, despite the application of skin surface sterilization. Therefore, we asked how similar in general the microbiome signature is for different sample types by comparing the microbial species detected in these samples. We first performed Non-metric Multidimensional Scaling (NMDS) analysis and principal coordinates analysis (PCoA), and then characterized the beta diversity of CSF and other specimen types using Bray-Curtis distances, a metric commonly used to evaluate microbiome difference among samples supported by Wilcoxon statistical significance. NMDS, PCoA (Supplementary Figure 2), and beta-diversity analysis revealed an overall clear separation of microbial communities for each sample type, except that microbiome in CSF_DNA specimens overlapped partially with negative controls (Figure 2a). Statistical analysis suggested beta-diversity between CSF_DNA and other sample types was significantly different from CSF_DNA self-comparison. However, there was no significant difference between CSF_DNA self beta-diversity and CSF_DNA-negative beta diversity (Wilcox test: p=0.59) (Figure 2b). In addition, the low diversity suggested the microbial communities in CSF_DNA and negative controls were highly similar. In fact, shared microbial taxa between CSF_DNA and negative control accounted for 42% and 22% of CSF_DNA and negative control, respectively. By contrast, 58% microbial taxa in CSF-DNA were detected in skin samples, whereas only 4% of skin microbes were found in CSF-DNA specimens. On one hand, these results indicated the microbial cells or DNA detected in CSF samples may partly have come from negative controls during sample collection. On the other hand, the high beta-diversity between skin and CSF specimens implied that the skin surface sterilization before lumbar punctures effectively prevented the contamination of CSF samples with most, if not all, skin microbes.

**Figure 2.**
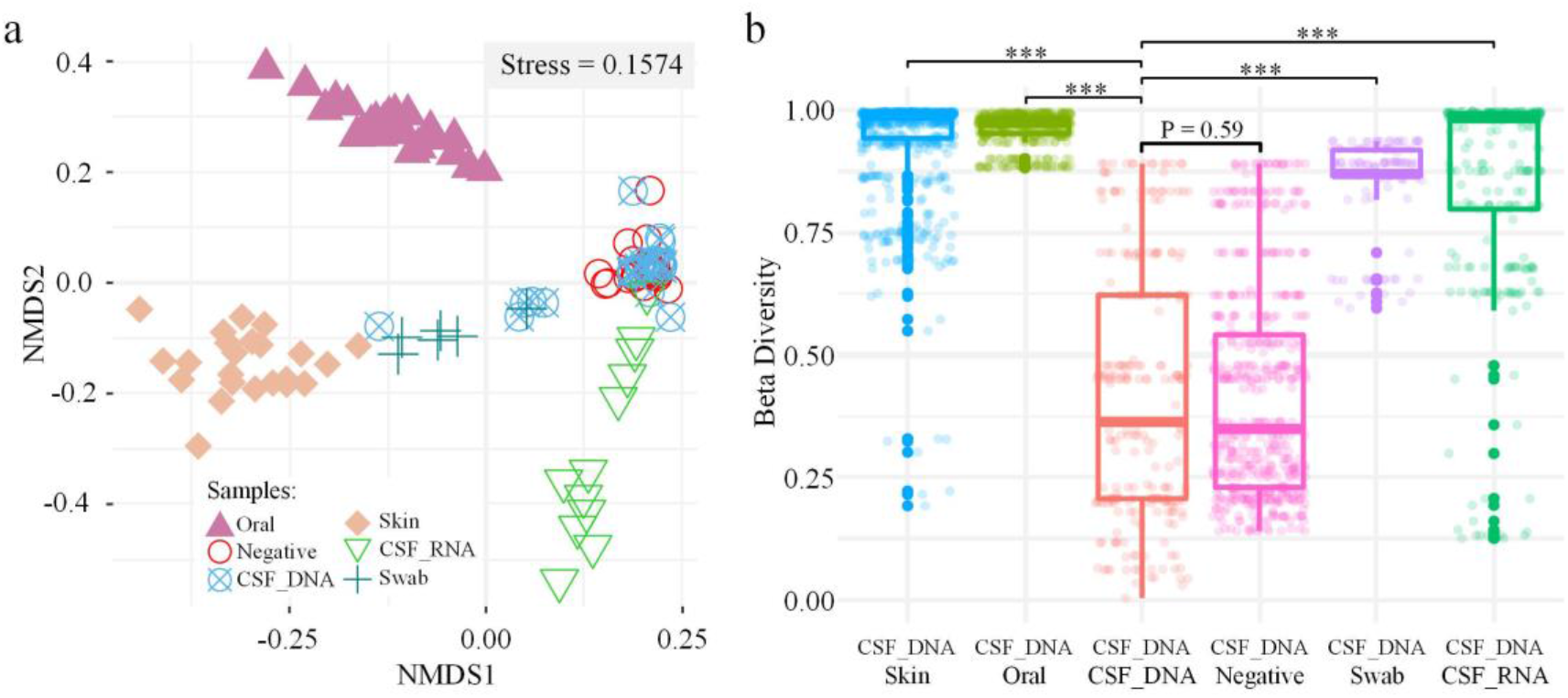
Microbiome similarity among sample types. (a) NMDS (Non-metric Multidimensional Scaling) analysis of microbial species detected from different sample types. Shapes and colors represent sample types. (b) Boxplot summarizing the beta diversity within CSF_DNA and between CSF_DNA and other specimens using Bray-Cruits dissimilarity. Statistical significance was assessed by Wilcoxon test whose significance level is indicated with asterisks (***: P<0.001).

### No microbiome is present in the CSF after subtracting microbes from controls

With the detected microbiome in CSF samples, we questioned whether these microbes were truly CSF inhabitants or simply brought in from external sources such as skins, negative controls and DNA extraction buffer. To verify whether the CSF contains de facto colonized microbial communities, we substracted the microbes collectively detected in negative control samples and DNA extraction buffer samples from microbes of each CSF_DNA sample, a method commonly used and previously described by human microbiome study(17). After substraction, 12 CSF samples contained no microbe, whereas the other 11 CSF samples contain a total of 14 microbial taxa including 11 viruses, 2 bacteria and 1 eukaryota. Since an introduction of microbes from skins could not be completely ruled out, we further checked whether the 14 taxa were present in skins as well and found that 6 of the 14 taxa were also found in the matching skin microbiome. This left eight microbial taxa after substraction as potential CSF inhabitating microbes, including five viruses (“*Bovine alphaherpesvirus 1*”, “*Escherichia virus V5*”, “*Klebsiella virus KP27*”, “*Macaca mulatta polyomavirus 1*”, “*Trichoplusia_ni_single_nucleopolyhedrovirus*”), two bacteria (*Hydrogenophilus thermoluteolus*, *Tepidimonas fonticaldi*), and one eukaryota (*Aspergillus turcosus*).

The detection of microbes using the metagenomic approach offers a glimpse of microorganisms present in certain niches. However, it remains uncertain whether these microbes are live or dead, as DNA from dead cells are also detectable by mNGS. Therefore, we further evaluated the physiological activities of the potential CSF-inhabitating microbes using metatranscriptomic sequencing, because the number of microbes detected by both in metagenomic and metatranscriptome would indicate active microbes may be present in CSF samples. CSF transcriptomics revealed transcripts for several microbial taxa including “*Equine infectious anemia virus*” and “*Cyprinid herpesvirus 3*” appearing in all samples, and *Escherichia_coli* and *Dasheen mosaic virus* appearing in eleven and nine samples, respectively. We then asked, for the eleven CSF-DNA samples with microbes left after substraction by negative controls and DNA extraction buffers, whether these microbes have detected in metatranscriptomic data. The result showed that only “*human alphaherpesvirus 1*” had signals from both CSF genomics and transcriptomics. However, “*human alphaherpesvirus 1*” also appeared in the skin, suggesting no active microbiome was detectable in CSF after removing this species potentially originated from skins. Although metagenomic analysis detected the one *Aspergillus turcosus* species from four individuals (Figure 3), no transcripts of *Aspergillus turcosus* were detected in metatranscriptomic, suggesting a lack of living cell activity. *Aspergillus turcosus* is well known as opportunistic human pathogens and can cause infections in individuals of compromised immune systems(32). How this fungal species (cells or DNA) reaches the CSF of the five healthy individuals is unknown, but it shows CSF, though without an active microbiome, might not be entirely free of opportunistic fungi which could potentially cause infections in central nervous systems when host immune system is compromised. Taken together, our study found no strong evidence supporting actively transcribed microbiome in the CSF.

**Figure 3.**
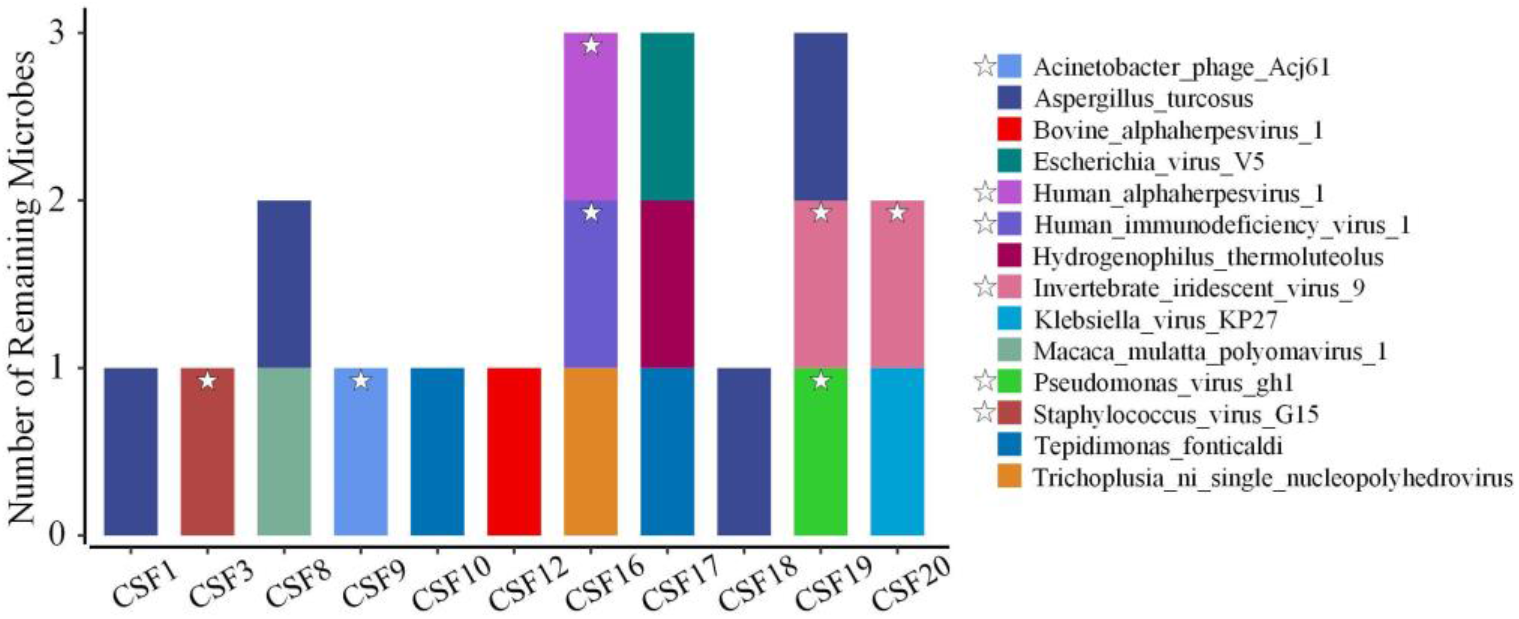
Microbes remained in the cerebrospinal fluids. Subtracting the microbes appeared in the negative control and DNA extraction buffer, 14 species (6 species labeled with star appeared in skin samples) remined in CSF genomic samples.

## DISCUSSION

Hereby, CSF samples from a cohort of 23 healthy individuals without neurological disease with a matched set of controls were collected for microbiome detection using culture-independent approach by a whole genome shotgun sequencing. The metagenomic data analysis indicated that there was no significant difference between CSF specimens and negative controls in beta diversity of detected microbes. In addition, no clear signal of active microbiome in the CSF samples was found by comparing CSF and contamination controls. Except *Aspergillus turcosus* appeared in four samples, no microbiome was present in more than two CSF samples after being subtracted by microbes in negative controls and DNA extraction buffer.

Compared with bacteria, more viruses were detected in CSF specimens. These viruses are mainly bacteriophages, most of which are also present in negative and skin samples. Although bacteriophages in the CSF have been reported, clear evidence for regular colonization of CSF by these viruses is lacking (19). Whether these viruses appear accidentally or colonized in CSF needs further investigation.

In our results, four CSF samples contained *Aspergillus* DNA, but no *Aspergillus* nucleic acid was detected in the corresponding RNA samples, suggesting that these *Aspergillus* DNA fragments may have come from contaminations. Due to the high sensitivity of mNGS, it can also detect trace amount of nucleic acid fragments released from dead microorganisms present in human periphery blood or tissues, experiment reagents and consumables. Furthermore, when using puncture to collect CSF specimen, tissues such as skin, muscle, blood vessels are potential sources of contamination. Except for strict disinfection measures before operation, constructing a database of colonizing microorganisms of these healthy tissues will enable subsequent bioinformatics analysis to filter out noise signals and reduce the false positive rate.

Highly sensitive mNGS represents a powerful tool for detecting microbiome at species resolution, especially for microbiome studies in specimens of low-abundance biomass, such as CSF. Currently, mNGS has become an important auxiliary method for clinical pathogenic diagnosis and treatment of infectious diseases. Main challenges of studying this issue have been an overall lack of CSF samples from healthy human subjects and the technically sound sampling as well as data analysis methods based on different reference databases and taxonomic strategies.

We focused on determining whether a CSF microbiome is present in healthy individuals without neurological disorders, a long-disputed issue in scientific and clinical research field. Our data analysis demonstrated that the microbiome of CSF was indistinguishable from contamination controls. It is intriguing but remains unclear whether a microbiome is present in CSF of patients diagnosed with diseases such as Alzheimer’s disease, multiple sclerosis, Parkinson’s disease and what roles the CSF microbiome plays in the development of these disorders.

In conclusion, using metagenomic combined with metatranscriptomic deep sequencing, we found microbiome profile in CSF samples was indistinguishable from that in contamination controls. Our data indicated that by current approaches there was no evidence to support the existence of a CSF microbiome in the populations without known neurological disorders. Such findings shall have great implications to human health especially neurological disorders and infections, providing a guide for disease diagnostics, prevention and therapeutics in clinical settings.

## MATERIALS AND METHODS

### Subjects

Twenty-three donors were recruited from the Xijing Hospital of the Fourth Military Medical University. All subjects were enrolled from obstetrics department in which the pregnant woman aged 23–40 years need intraspinal anesthesia before the caesarean section. Subjects who have suffered from central nervous system infection disease (eg, meningitis, encephalitis) or any systemic infection disease and autoimmune disease (eg, hepatitis, tuberculosis, systemic lupus erythematosus, rheumatism) and have received antibiotics treatment in the past six months prior to sample collection were excluded. We also excluded subjects with a history of hypertension, diabetes, heart disease, cancer and neurological disease (eg, Alzheimer’s disease, Parkinson’s disease, multiple sclerosis, epilepsy).

### Sample collection

Lumbar puncture was performed in the 23 subjects enrolled in this study and the CSF samples were collected in a 4ml centrifugal tube with syringe and then stored in a −80 °C freezer for metagenomics analysis. Twelve CSF samples were randomly selected from 23 pregnant women for metatranscriptome studies and RNA protection reagent was added to the CSF immediately after collection. Then, the samples were centrifuged and the pellets stored at −80 °C for metatranscriptomic sequencing. Meanwhile, normal saline was collected with syringes for environmental controls (negative control). Furthermore, oral and skin samples were selected from 23 enrolled subjects as one-to-one matched positive control. For skin positive controls: The back skin of 5×5 cm^2^ areas around the puncture site (L3-L4 intervertebral space) were swabbed with a sterile cotton swab before skin clean with povidone iodine. To maximize microbial load, no bathing was permitted within 24 hours of sample collection. For oral positive controls, all subjects were forbidden to eat and drink six hours before operation. The surfaces of tongue, buccal fold, hard palate, soft palate, tooth, gingiva and saliva were swabbed with sterile swab. Unused sterile swabs were collected for negative controls (“sterile swab”). Details of Matching information between samples are described in Supplementary Table S2.

### DNA extraction and purification

DNA was isolated using the QIAamp DNA Mini Kit (Qiagen 51304) according to the manufacturer’s instructions. 1) DNA extraction from swabs: Swab tips were cut and placed in a 2 ml microcentrifuge tube and then 400 μl PBS were added. Next, 20 μl of proteinase K and 400 μl of buffer AL were added, vortexed for 10 s, and the solution was incubated for 15 min at 56 °C. And then added 400 μl ethanol (100%) and mixed again by vortexing. Lastly, DNA purification was performed with buffer AW1 and AW2 using QIAamp Mini spin column, followed by elution with 35 μl of buffer EB. 2) DNA extraction from CSF and normal saline controls: 200 μl sample was added into the microcentrifuge tube, and then added 20 μl of proteinase K and 200 μl of buffer AL respectively, vortexed for 10 s, and the solution was incubated for 15 min at 56 °C. Next, added 200 μl ethanol (100%) and mixed again by vortexing. DNA purification was performed as described above.

### Metagenomics library construction

For preparation of metagenomics libraries, the QIAseq FX DNA Library Kit (Qiagen; 180715) was used. The construction involved five steps: 1) Fragmentation and End-repair: to generate 200–300 bp fragments, 32.5μl purified DNA were fragmented by incubation with FX buffer 5μl, FX enhancer 2.5μl and 10 μl FX enzyme mix at cycling program: 4 °C 1minute→ 32 ° C 12minutes→ 65 ° C 30minutes→ 4 ° C hold. 2) Adapter ligation: 5 μl of adaptor, 20 μl of ligation buffer, 10 μl of DNA ligase and 15 μl of nuclease-free water were added and incubated for 15 minutes at 20 °C to initiate adapter ligation. Adapter ligation cleanup was performed immediately, 3) Adapter ligation cleanup: 80 μl of resuspended AMPure® XP beads (0.8×) were added to each ligated sample and mix well by pipetting. Next, the mixture was incubated for 5 minutes at room temperature and then the beads were pelleted on a magnetic stand (Invitrogen) for 2 minutes. The supernatant was discarded and the pallet was washed twice with 200 μl of 80% ethanol, then the beads were eluted with 52.5 μl of buffer EB. Subsequently, 50 μl of supernatant was transfered into a new 1.5 ml microcentrifuge tube and a second purification was performed with 50 μl (1×) AMPure® XP beads. The final, 23.5 μl of purified DNA sample was obtained. 4) Amplification of library DNA: 25 μl of HiFi PCR Master Mix, 1.5 μl of Primer Mix and 23.5 μl of library DNA were added in PCR tube. PCR enrichment was performed under the cycle conditions: 2 minutes at 94 °C, 12 × (20 s at 98 °C, 30 s at 60 °C, 30 s at 72 °C), and 1 minute at 72 °C. The final, to obtain libraries, the PCR products were purified with AMPure XP beads as described above.

### RNA extraction and purification

Total RNA was extracted using the RNeasy Mini Kit (Qiagen; 74104) according to the manufacturer’s instructions. The pellet of each sample which has been treated with RNA protection reagent as described above, was resuspended in 100μl TE buffer containing lysozyme, and Proteinase K was added into the mixture, then incubated for 10 minutes at room temperature. 350μl of buffer RLT was added and vortexed vigorously. The final, RNA isolation and purification was performed with buffer AW1 and RPE respectively using RNeasy Mini spin column, followed by elution with RNase-free water.

### RNA library preparation for metatranscriptomics sequencing

For construction of RNA libraries, the QIAseq FX Single Cell RNA Library Kit (Qiagen; 180733) was used. The construction involved five steps: 1) Genomic DNA (gDNA) removal: 8 μl of purified RNA and 3 μl of NA denaturation buffer were added into a sterile PCR tube and incubated for 3 minutes at 95 °C. To remove genomic DNA, 2 μl of gDNA wipeout buffer was added and incubated for 10 minutes at 42 °C. 2) Reverse transcription: 4 μl of RT/Polymerase buffer, 1 μl of random primer, 1 μl of Oligo dT primer and 1 μl of Quantiscript RT enzyme mix were added in each sample and reverse transcription was carried out for 60 minutes at 42 °C. 3) Ligation: 8 μl of ligase buffer and 2 μl of ligase mix were added into the RT reaction and incubated at 24°C for 30 minutes. 4) Whole transcriptome amplification: 1 μl of REPLI-g SensiPhi DNA Polymerase and 29μl of reaction buffer were used for Multiple Displacement Amplification (MDA), then incubate at 30°C for 2 h. The final, an approximate length of 2000–70,000 bp amplified cDNA was produced. 5) Enzymatic Fragmentation: The amplified cDNA was diluted 1:3 in H_2_O sc, 10 μl of the diluted DNA and FX Enzyme Mix were used to obtain 300 bp library fragment with reaction conditions: 4 °C 1minute→32°C 15minutes→65°C 30minutes→4°C hold. 6) Adapter ligation: 5μl of adapter and 45μl of ligation master mix were added into each sample and incubated at 20° C for 15 minutes. Subsequently, the adapter ligation cleanup was performed with AMPure XP beads as described above. The final, purified libraries were obtained ready for sequencing without further PCR amplification.

### Next generation equencing

Shotgun sequencing was performed on Illumina Hiseq platform for all samples (paired end library of 150-bp and 150-bp read length). Approximately, 25 Gb and 5 Gb of raw paired-end reads were obtained per sample in the CSF genomics samples and negative samples, respectively.

### Data quality control

To reduce the impact of host reads on the results, we need to remove human reads involved in the raw sequencing data before bioinformatics analysis. KneadData(29) (v0.7.2), a widely used tool, is designed to perform quality control on metagenomic and metatranscriptomic sequencing data, especially for microbiome experiments. All reads were filtered using KneadData with the following trimmomatic options: ILLUMINACLIP: TruSeq3-PE-2.fa:2:30:10:8:true, SLIDINGWINDOW:4:20, MINLEN:50 and bowtie2 options: --very-sensitive, --dovetail. The proportion of human reads in CSF genomics samples is up to 92%.

### Detecting potential microbiome

MetaPhlAn(30) (v 3.0.1) is a computational tool for profiling the composition of microbial communities (bacteria, archaea, viruses and eukaryotes) from shotgun sequencing data. Based on ~1.1M unique clade-specific marker genes identified from ~100,000 reference genomes, MetaPhlAn can profile unambiguous taxonomic assignments and accurate estimation of organismal relative abundance in species-level resolution. Classifying the reads to marker genes database, MetaPhlAn outputs a file containing detected microbes and relative abundance in different level. MetaPhlAn ran with custom parameters: --add_viruses--input_type fastq--read_min_len 50. It’s worth noting that MetaPhlAn (previous version 2) was the only bioinformatics tool with 0% false positive relative abundance and the best diversity estimate(33).

### β-diversity and phylogenetic analysis

Using R (version 3.6.3) with R studio environment, β-diversity (between-sample diversity) was estimated by Bray-Curtis dissimilarity in vegan package. All figures are ploted using R.

## Supplementary files

**Supplementary Table S1.** Microbes detected in different samples.

## Availability of data and materials

The clean sequence data reported in this paper have been deposited in the Genome Sequence Archive in BIG Data Center(34, 35), Chinese Academy of Sciences, under accession number PRJCA004977<u>XXXXX</u> that are publicly accessible at https:///bigd.big.ac.cn/bioprojectXXX.

## Acknowledgements

We thank Peng Jia, Tingjie Wang, Ningxin Dang, Honghui Shen and Tun Xu for helpful discussions regarding data analysis and Jing Hai for administrative and technical support. We thank the High-Performance Computing Cluster of the First Affiliated Hospital of Xi’an Jiaotong University for data processing.

This study was supported by the National Key R&D Program of China (Grant Nos. 2018YFC0910400, 2017YFC0907500, and 2016 YFC0904501), the National Natural Science Foundation of China (Grant Nos. 31671372, 61702406, 31701739, and 31970317), the National Science and Technology Major Project of China (Grant No. 2018ZX10302205), as well as the General Financial Grant from the China Postdoctoral Science Foundation (Grant Nos. 2017M623178 and 2017M623188).

## Ethics Statement

This study was approved by the Ethics Committee of the Xijing Hospital of the Fourth Military Medical University. All procedures were conducted in accordance with the approved guidelines. All patients read and signed the consent form before sample collection.

---

**Supplementary Figure S1:**
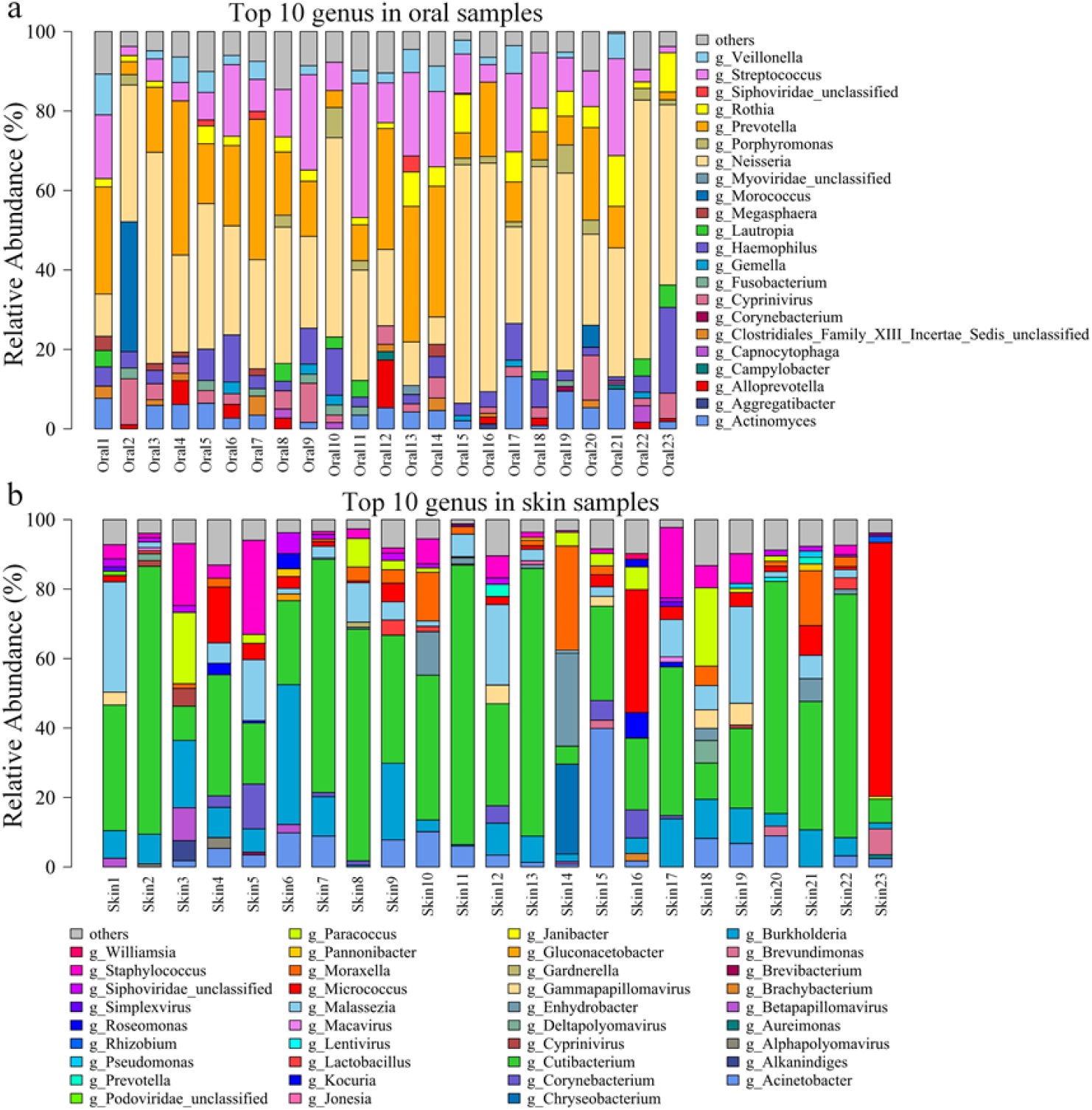
Top 10 genus in oral and skin samples, respectively. Microbial community structures of 23 Oral (figure S1a) and Skin (figure S1b) samples shown in a stacked barplot that summarizes the relative abundance of different genus detected.

**Supplementary Figure S2:**
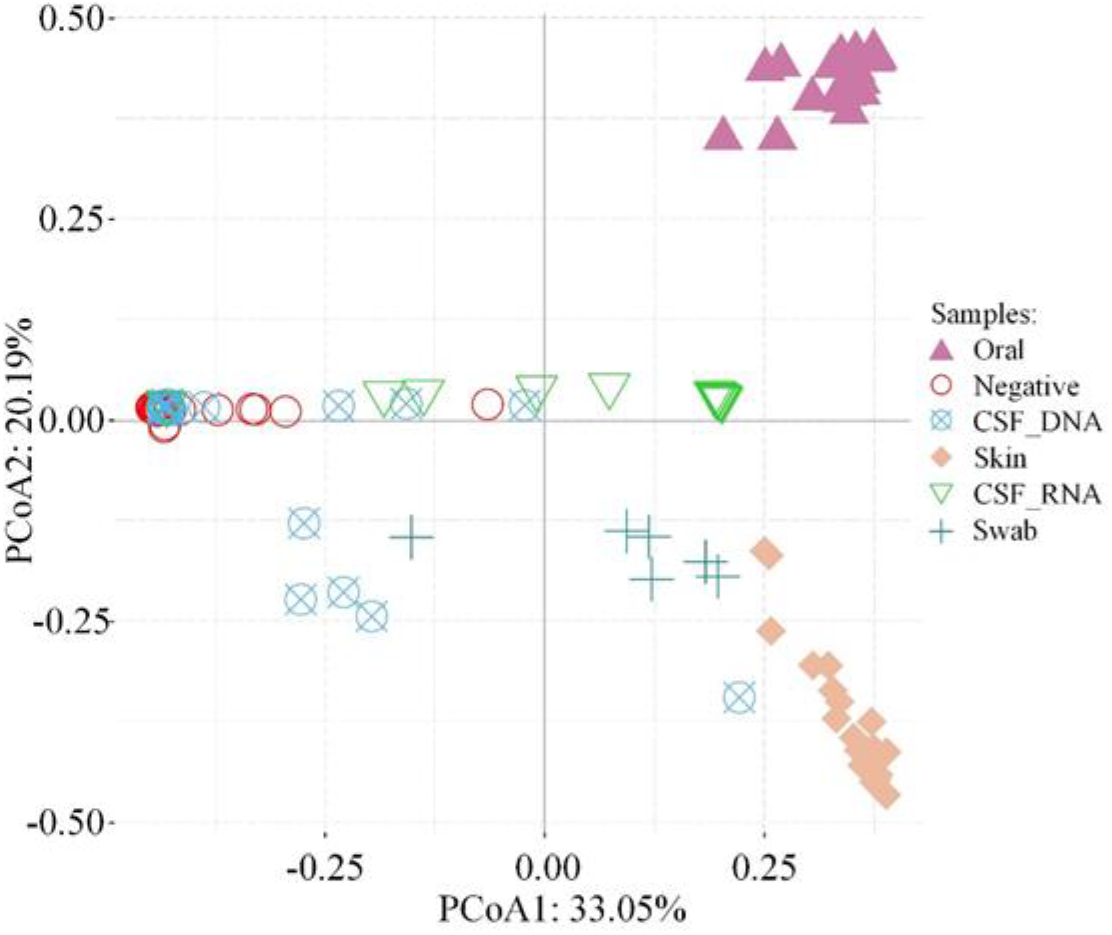
Principal Coordinates Analysis (PCoA) analysis of microbial species detected from different sample types. PCoA (Principal Coordinates Analysis) analysis of microbial species detected from different sample types. Shapes and colors represent sample types.

**Supplementary Table S2:** Sample labels and matching information.

| Table S1: Samples label and matching information |  |  |  |  |  |  |  |
| --- | --- | --- | --- | --- | --- | --- | --- |
| CSF_DNA | Negative | Skin | Oral | CSF_RNA | Swab | RNA_buffer | DNA_buffer |
| 1 | 1 | 1 | 1 | 1 | 1 | 1 | 1 |
| 2 | 2 | 2 | 2 |  | 2 | 2 | 2 |
| 3 | 3 | 3 | 3 |  | 3 | 3 | 3 |
| 4 | 4 | 4 | 4 | 4 | 4 |  |  |
| 5 | 5 | 5 | 5 |  | 5 |  |  |
| 6 | 6 | 6 | 6 |  | 6 |  |  |
| 7 | 7 | 7 | 7 |  |  |  |  |
| 8 | 8 | 8 | 8 | 8 |  |  |  |
| 9 | 9 | 9 | 9 | 9 |  |  |  |
| 10 | 10 | 10 | 10 | 10 |  |  |  |
| 11 | 11 | 11 | 11 |  |  |  |  |
| 12 | 12 | 12 | 12 |  |  |  |  |
| 13 | 13 | 13 | 13 |  |  |  |  |
| 14 | 14 | 14 | 14 | 14 |  |  |  |
| 15 | 15 | 15 | 15 | 15 |  |  |  |
| 16 | 16 | 16 | 16 | 16 |  |  |  |
| 17 | 17 | 17 | 17 |  |  |  |  |
| 18 | 18 | 18 | 18 | 18 |  |  |  |
| 19 | 19 | 19 | 19 | 19 |  |  |  |
| 20 | 20 | 20 | 20 |  |  |  |  |
| 21 | 21 | 21 | 21 | 21 |  |  |  |
| 22 | 22 | 22 | 22 |  |  |  |  |
| 23 | 23 | 23 | 23 | 23 |  |  |  |
| CSF_DNA: Cerebrospinal fluids metagenomic |  |  |  |  |  |  |  |
| CSF_RNA: Cerebrospinal fluids metatranscriptomic |  |  |  |  |  |  |  |

